# FastMLST: A multi-core tool for multilocus sequence typing of draft genome assemblies

**DOI:** 10.1101/2020.10.13.338517

**Authors:** Enzo Guerrero-Araya, Marina Muñoz, César Rodríguez, Daniel Paredes-Sabja

**Author notes:** To whom correspondence should be sent **Contact:** Enzo Guerrero-Araya and Daniel Paredes-Sabja.

## Abstract

Multilocus Sequence Typing (MLST) is a precise microbial typing approach at the intra-species level for epidemiological and evolutionary purposes. It operates by assigning a sequence type (ST) identifier to each specimen, based on a combination of allelic sequences obtained for multiple housekeeping genes included in a defined scheme. The use of MLST has multiplied due to the availability of large numbers of genomic sequences and epidemiological data in public repositories. However, data processing speed has become problematic due to datasets’ massive size. Here, we present FastMLST, a tool that is designed to perform PubMLST searches using BLASTn and a divide-and-conquer approach. Compared with mlst, CGE/MLST, MLSTar, and PubMLST, FastMLST takes advantage of current multi-core computers to simultaneously type thousands of genome assemblies in minutes, reducing processing times by at least 4-fold and with more than 99.95% consistency.

**Availability and Implementation:** The source code, installation instructions and documentation are available at https://github.com/EnzoAndree/FastMLST

## 1. Introduction

Multilocus Sequence Typing was a revolutionary attempt to standardize the molecular typing of *Neisseria meningitidis* (Maiden, et al., 1998) that has been replicated in multiple pathogens of global relevance (Dingle, et al., 2001; Griffiths, et al., 2010; Martin-Rodriguez, et al., 2019; Meats, et al., 2003). This technique assigns a bacterial isolate a sequence type (ST) based on a combination of alleles from an optimal set of housekeeping genes defined for each species. Even though there are methods with higher discrimination power (cgMLST/wgMLST), MLST offers several advantages, such as (1) a common language for typing, (2) simple implementation, and (3) a vast amount of information collected throughout history (Kimura, 2018).

Currently available programs for ST determination from assembled genomes i.e. mlst (Seemann, 2015), CGE/MLST (Larsen, et al., 2012), MLSTar (Ferrés and Iraola, 2018), and the online tool PubMLST (Jolley, et al., 2018) deliver results in 1-9 minutes when dealing with hundreds of genomes. However, since most of them do not (or inefficiently) implement parallel computing, processing of tens of thousands of genomes can be extremely time-consuming. In addition, their outputs are typically restricted to a table with allelic profiles and do not deliver a file with concatenated allele sequences necessary for subsequent phylogenetic analyses (Glaeser and Kampfer, 2015).

In response to these limitations, we developed FastMLST, a tool focused on parallel computing via a divide-and-conquer approach (Smith, 1985) that performs BLASTn searches (Camacho, et al., 2009) against the sequences of MLST schemes deposited in the PubMLST database (Jolley, et al., 2018). FastMLST delivers two main outputs: a table the allelic profiles detected for a genome query and a multi-FASTA file containing their sequences.

## 2. Methods

### 2.1. Implementation

FastMLST is an open source standalone software implemented in Python 3 that only requires BLASTn as an external program. FastMLST is easy to install via the Anaconda Package Manager (https://anaconda.org/bioconda/fastmlst) and is in constant development.

### 2.2. Key analysis steps

#### PubMLST database setup

FastMLST can retrieve and update the PubMLST database (Jolley, et al., 2018) with its ‘--update- mlst’ option. It automatically processes the 150 currently available schemes (October 2020) and prepares the system for subsequent deployment.

#### Allele searches

FastMLST is based on a divide-and-conquer approach, whereby each genome is processed in parallel. Allele searches are performed with BLASTn using the public, regularly updated, and centralized database maintained at PubMLST (Jolley, et al., 2018).

#### Scheme identification

The MLST scheme to use for each query genome is inferred by an automatic scoring system, similar to the one reported by (Seemann, 2015). This system assigns known, full-length alleles a maximum score of 100 points, 70% of the score to alleles of the expected length but with SNPs (95% identity by default), or 20% of the score to alleles with unexpected lengths. At the end, the MLST scheme with the highest overall score is selected for further analyses. Alternatively, it is possible to choose among schemes manually. When multiple perfect hits to a single scheme are found, FastMLST reports all hits to alert the user of potential contaminations.

## 3. Performance benchmark

We assessed the capability of FastMLST to identify STs correctly for three bacterial species with unequal numbers of scheme alleles. To this end, we measured the concordance over ST assignment between FastMLST and PubMLST (Jolley, et al., 2018). The species tested were *Cutibacterium acnes, Enterococcus faecium*, and *Streptococcus pneumoniae*, which as of September 11^th^ 2020 included 153, 1,833 and 16,301 STs, respectively. Besides, we compared the processing speeds of FastMLST and four others widely used software: mlst v2.11, CGE/MLST v2.0.4, MLSTar v1.0, and the online tool PubMLST.

A total of 278 genomes for *C. acnes*, 1,915 for *E. faecium*, and 7,996 for *S. pneumoniae* obtained from GenBank using ncbi-genome-download v0.2.11 (https://github.com/kblin/ncbi-genome-download) were used in the analysis of ST assignment concordance (**Supplementary Table 1**). Compared with PubMLST result (**Supplementary Tables 2, 3 and 4**), we show that FastMLST (**Supplementary Tables 5, 6 and 7**) reaches a concordance of 99.96% (n=9,502/9,506) (Table 1). As to the inconsistencies detected, FastMLST failed to assign STs to 4 genomes with multiple perfect hits against the same locus, indicating that they are possibly contaminated (Supplementary Table 6 and 7). Additionally, FastMLST reported 401 STs that remained undetected by PubMLST (7 in *C. acnes*, 216 in *E. faecium*, and 178 in *S. pneumoniae*) (Supplementary Table 5, 6 and 7). These new ST are derived from new alleles or new combinations of known alleles (Table 1). On the other hand, FastMLST missed some full-length alleles due to incomplete coverage (<100%) (Table 1).

**Table 1.** Concordance analysis between FastMLST and PubMLST using 10,189 genomes.

| Dataset | Genomes | %GC | ST assigned by PubMLST<sup>a</sup> | ST assigned by FastMLST<sup>b</sup> | Concordance<sup>c</sup> | New ST with full-length alleles<sup>d</sup> | Missing alleles (<100% coverage)<sup>e</sup> |
| --- | --- | --- | --- | --- | --- | --- | --- |
| *C. acnes* | 278 | 60% | 259 | 259 | 259/259 (100%) | 7 | 12 |
| *E. faecium* | 1915 | 38% | 1545 | 1544 | 1544/1545 (99.94%) | 216 | 155 |
| *S. pneumoniae* | 7996 | 39% | 7702 | 7699 | 7699/7702 (99.96%) | 178 | 119 |

<sup>a</sup> A genome was included in this category if an ST was assigned using PubMLST.
<sup>b</sup> A genome was included in this category if an ST was assigned using FastMLST.
<sup>c</sup> A ST assignment was considered concordant if the ST assigned by PubMLST and FastMLST was identical.
<sup>d</sup> A genome with novel alleles (100% coverage). It is a designation only present in FastMLST.
<sup>e</sup> Details of genomes with missing alleles detected using FastMLST.

The speed of FastMLST, MLSTar, PubMLST and mlst (expressed in genomes/minute) was compared in a workstation equipped with 2 Intel(R) Xeon(R) CPU E5-2683 v4 @ 2.10GHz using 1, 2, 4, 8, 16, 32 or 64 cores. Each program was run in triplicate with a sample of 278 *C. acnes* genomes each time. The results for PubMLST were obtained remotely because it does not offer a standalone version. When a single core was used, mlst v2.11 achieved the best performance, with 93 genomes/minute (Figure 1A). However, mlst v2.11 did not speed up when its multiple processing option was enabled (Figure 1A). MLSTar can also use multiple processors, but it is at least 4-fold slower than FastMLST (Figure 1A and 1B). FastMLST’s processing speed grew exponentially as more cores were deployed (Figure 1B), reaching a speed of 1,642 genomes/minute using 64 cores. This allows processing of 16,000 genomes in less than 10 minutes.

**Figure 1.**
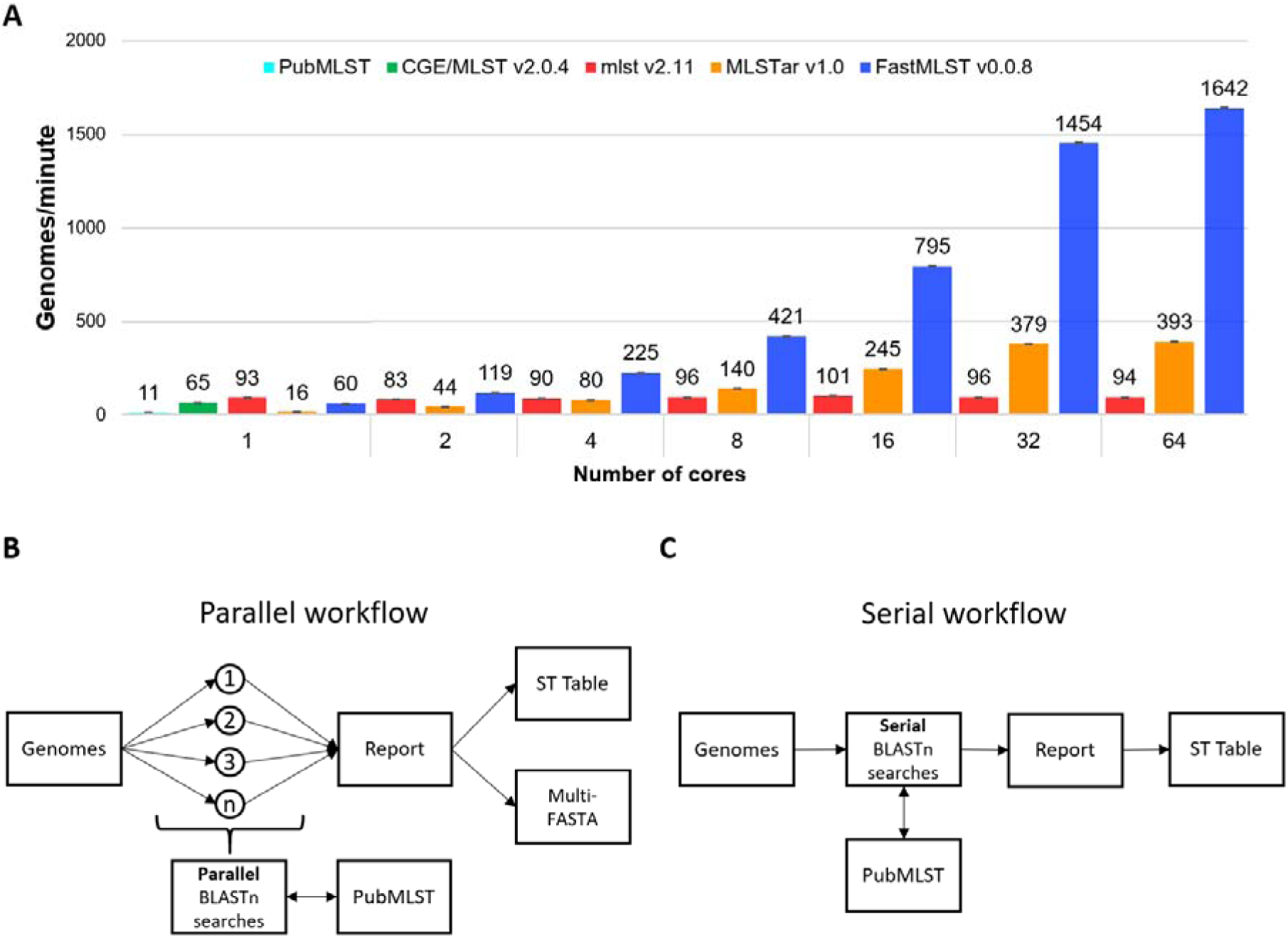
Processing speed and simplified comparative-workflow. **(A)** Processing speed (genomes/minute) of FastMLST and standard software for MLST determination (mlst, PubMLST, CGE/MLST and MLSTar) was compared by triplicate using a dataset composed of 278 C. acnes genomes and 1, 2, 4, 8, 16, 32, and 64 cores. Error bars represent the standard error of three replicates. Only mlst, MLSTar and FastMLST natively support parallel processing. **(B)** FastMLST and MLSTar workflow. **(C)** mlst, PubMLST and CGE/MLST workflow.

## 4. Final remarks

FastMLST assigns STs to thousands of genomes in minutes with the additional plus of generating a multi-FASTA file with concatenated allele sequences suitable for downstream phylogenetic analysis. These unique features have the potential to boost future research.

## Funding

This work was supported by a Doctoral fellowship 21181536 from ANID to E.G-A., and by EU-Lac project “Genomic Epidemiology of *Clostridium difficile* in Latin America (T020076)”, ANID – Millennium Science Initiative Program – NCN17_093, and Start-up funds of Texas A&M University to D.P-S.

## Notes

### Competing Interest Statement

The authors have declared no competing interest.

